# Inhibition of NF-κB Signaling by the Reactive Glycolytic Metabolite Methylglyoxal

**DOI:** 10.1101/2025.05.13.653579

**Authors:** Caroline Stanton, Woojin Choi, R. Luke Wiseman, Michael J. Bollong

**Affiliations:** Department of Chemistry, The Scripps Research Institute, La Jolla, CA, 92037, USA; Department of Molecular and Cellular Biology, The Scripps Research Institute, La Jolla, CA, 92037, USA

## Abstract

The NF-κB family of transcription factor complexes are central regulators of inflammation, and their dysregulation contributes to the pathology of multiple inflammatory disease conditions. Accordingly, identifying pharmacological mechanisms that restrain NF-κB overactivation remains an area of key importance. Here, we demonstrate that inhibition of the glycolytic enzyme phosphoglycerate kinase 1 (PGK1) with the small molecule inhibitor CBR-470-2 results in attenuated NF-κB signaling, decreasing transcriptional output in response to several canonical NF-κB activating stimuli. Mechanistically, PGK1 inhibition promotes the accumulation of the glycolytic metabolite methylglyoxal, which crosslinks and inactivates NF-κB proteins, limiting the phosphorylation and nuclear translocation of these transcription factor complexes. This work establishes a key connection between central carbon metabolism and immune signaling and further supports the notion that PGK1 inhibition may be a viable strategy to increase cellular survival and dampen inflammation in disease.

## Introduction

The family of Nuclear factor-kappa B (NF-κB) transcription factor complexes regulate a diverse array of cellular processes including inflammation, growth, differentiation, and survival^1^. Each NF-κB transcription factor complex is typically nucleated by one of two so-called Class 1 proteins, NF-κB1 (also called p50) and NF-κB2 (also called p52), and one of three Class 2 proteins, RelA (also called P65), RelB, and c-Rel. Both Class 1 and 2 proteins contain a conserved Rel Homology domain (RHD)^2^, which mediates formation of homo- or hetero-dimers between these proteins and their ability to bind regulatory DNA response elements. Class 2 proteins also contain a transcriptional activation domain, and, as such, Class 1 proteins rely on the association with one of these factors to elicit transcription^3^. Among the potential NF-κB family complexes that can form, the P50/RelA heterodimer is the most characterized in the literature^3,4^.

Under basal conditions, NF-κB complexes are sequestered in the cytosol through interactions with the IκB family of inhibitory proteins, including IκBα^5^. Canonical NF-κB signaling is triggered by a variety of signals including ligands of pattern recognition receptors like TLR4 (e.g., lipopolysaccharide), TNF receptor superfamily members (e.g., TNF-a), cytokine receptors, B-cell receptors, and T-cell receptors^6^. Positive signaling through these pathways all result in the phosphorylation-dependent activation of the multi-subunit IκB kinase (IKK) complex, consisting of the catalytic subunits IKKα and IKKβ and the regulatory subunit NF-κB essential modulator (NEMO)^7^. The active IKK complex then subsequently phosphorylates two N-terminal serine residues of IκB, triggering its ubiquitin-dependent proteasomal degradation and the concomitant release of NF-κB dimers to translocate to the nucleus where they activate transcription at target loci^8^.

NF-κB activity must be tightly regulated to provide essential cellular growth cues without inducing unnecessary and potentially damaging pro-inflammatory signaling. Among the many factors which activate an inflammatory NF-κB-driven transcriptional response are reactive oxygen species (ROS), as NF-κB was the first redox responsive transcription factors described in literature^9,10^. NF-κB signaling is activated by a variety of ROS, leading to the expression of pro-inflammatory cytokines to stimulate immune cell-driven resolution of tissue damage^11^. Acting in opposition to NF-κB is the oxidative stress responsive transcription factor NRF2 (NFE2L2), which senses ROS and electrophilic chemicals through several sensor cysteines in its cytoplasmic repressor protein KEAP1 (Kelch-like ECH-associated protein 1), and, when activated promotes the resolution of inflammation, the inactivation of NF-κB, and the dampening of innate immune signaling^12,13^. Given the intimate interplay between these two signaling systems, it is perhaps not surprising that a number of pro-protective electrophilic small molecules, including polyphenols, triterpenoids, and others have been characterized both as NRF2 activators (by covalent inactivation of KEAP1), as well as NF-κB inhibitors (by modifying several reactive cysteines IKK proteins as well as p65 and p50^14-18^). Notably, two endogenously occurring electrophilic metabolites, itaconate and fumarate, as well as their cell penetrant esters, have also been shown to both activate NRF2 and inactivate NF-κB through covalent modification-based mechanisms, raising the intriguing possibility that certain endogenous metabolites, in addition to their metabolic roles, may provide sentinel signaling cues to regulate essential processes related to cellular survival^19-23^.

We previously reported the discovery of the CBR-470 series of compounds, small molecules which inhibit the enzymatic activity of the glycolytic enzyme phosphoglycerate kinase 1 (PGK1)^24-27^. PGK1 inhibition leads to the buildup of upstream triose phosphates and their degradation product, the reactive metabolite methylglyoxal (MGO). MGO can covalently modify reactive cysteines as well as form non-enzymatically derived crosslinking MICA (methyl imidazole crosslink between cysteine and arginine)-based modifications between proteins. We found MGO to crosslink and inactivate KEAP1^24^, as well as NLRP3 (NOD-, LRR- and pyrin domain-containing protein 3), a pattern recognition receptor which regulates cleavage of pro-inflammatory cytokines via formation of the inflammasome^28^.

Given the substantial crosstalk between KEAP1, NLRP3, and NF-κB, we hypothesized that MGO might modulate NF-κB signaling in some way. Here, we show that that PGK1 inhibition by CBR-470-2 additionally inhibits NF-κB, discouraging transcriptional activation in response to diverse activating stimuli. Augmented levels of MGO result in the covalent crosslinking and inactivation of multiple central signaling proteins in the pathway, inhibiting their capacity to relay phosphorylation.

## Results and Discussion

### NF-κB signaling is inhibited by pre-treatment with PGK1 inhibitors

Given the capacity of numerous KEAP1 modifying electrophiles to inactivate NF-κB, we first sought to determine if the previously reported small molecule inhibitor of phosphoglycerate glycerate kinase 1 (PGK1), CBR-470-2, which results in a disruption of glycolytic flux and an accumulation of the reactive metabolite methylglyoxal (MGO), impacts the activity of NF-kB. Using the human monocytic leukemia derived THP-1 Dual reporter cell line which expresses secreted alkaline phosphatase (SEAP) under the control of an NF-κB binding element (a minimal IFN-B promoter fused to five copies of the NF-κB consensus transcriptional response element and three copies of the c-Rel binding site), we demonstrated that CBR-470-2 dose dependently inhibits NF-κB activity induced by both lipopolysaccharide (LPS) and TNF-α, which signal through different upstream receptors (TLR4 and TNFR respectively) (**Fig. 1A, B**). A similar inhibitory effect was observed using a human adenocarcinoma derived A549 Dual reporter cell line expressing secreted alkaline phosphatase (SEAP) under the control of the same NF-κB DNA binding element (**Fig. S1A, B**). We then confirmed that this inhibitory effect was also observed in wildtype THP1 cells by measuring transcript levels of the NF-κB target genes *IL1B, TNF,* and *IL6* as assessed by RT-qPCR after stimulation by LPS or TNF-α (**Fig. 1C-E, Fig. S1C, D**). CBR-470-2 was also found to decrease the nuclear localization of the P65 subunit of NF-κB following TNF-α stimulation in high content imaging-based immunofluorescence experiments (**Fig. 1F, G**), collectively demonstrating that CBR-470-2 inhibits the localization and transcriptional output of NF-κB.

**Figure 1.**
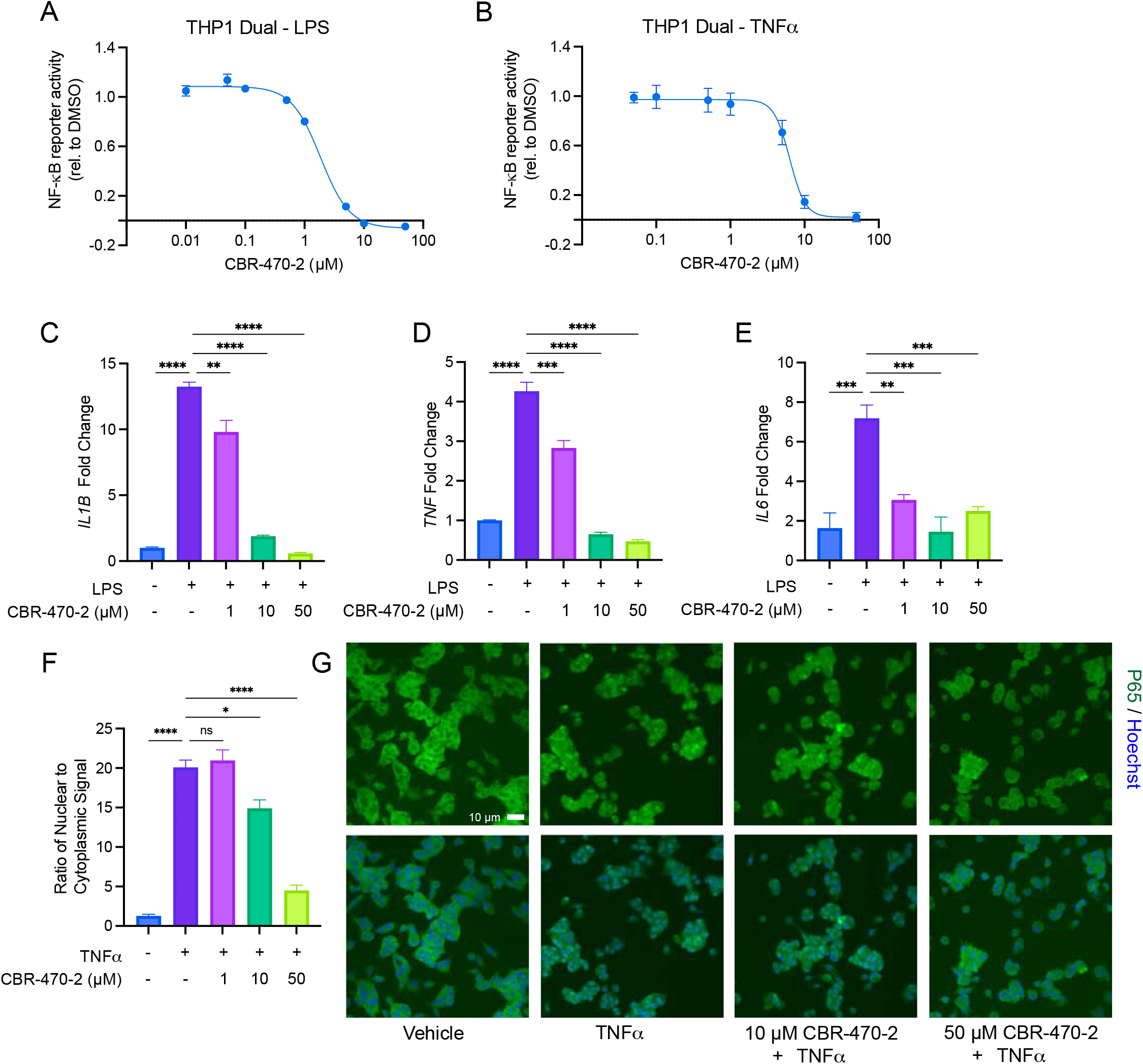
CBR-470-2 inhibits NF-κB signaling. (A, B) Relative NF-κB transcriptional activity as measured by SEAP reporter in THP-Dual cells pretreated with CBR-470-2 for 6 h and then stimulated with LPS (A) or TNF-α (B) overnight. Error bars show SEM for n = 5 replicates (A) or n = 4 replicates (B). (C-E) Relative transcript level of *IL1B* (C), *TNF* (D), or *IL6* (E) from THP1 cells pretreated with CBR-470-2 for 6 h and then stimulated with LPS overnight. Error bars show SEM for n = 3 replicates. **P<0.01, ***P<0.001, \*\*\*\**P*<0.0001 for ordinary one-way analysis of variance (ANOVA) with Tukey correction for multiple comparisons between conditions. (F) Ratio of nuclear to cytosolic localization of RelA in A549 Dual cells pretreated with CBR-470-2 for 6 h and stimulated with TNF-α for 1 h. Error bars show SEM for n = 3 replicates. *P<0.05, \*\*\*\**P*<0.0001 for ordinary one-way analysis of variance (ANOVA) with Tukey correction for multiple comparisons between conditions. (G) Representative images for nuclear localization of RelA from 1F.

### CBR-470-2 inhibits NF-κB through a covalent mechanism involving PGK1 inhibition

We then sought to identify the mechanism by which CBR-470-2 inhibits NF-κB. We initially characterized CBR-470-2 as a NRF2 activator. Because NRF2 activation is known to inhibit NF-κB, we next evaluated if NRF2 was relevant to the mechanism by which CBR-470-2 inhibits NF-κB. From experiments involving the co-treatment of THP-1 Dual cells with the NRF2 inhibitor ML385 and CBR-470-2, we observed no reduction in the potency or efficacy by which CBR-470-2 inhibited NF-κB reporter output in THP-1 Dual cells, suggesting that NRF2 activation is not relevant to the observed inhibitory mechanism (**Fig. 2A**). We then demonstrated that pretreatment of THP1 Dual and A549 Dual cells with 1 mM GSH, which quenches reactive electrophiles including MGO, markedly decreased the potency with which CBR-470-2 inhibited NF-κB reporter output induced by both TNF-α and LPS stimulation, suggesting that CBR-470-2 might be acting through a mechanism involving the augmentation of an reactive electrophile (**Fig. 2B, C, Fig. S2A-C**). CBR-470-2 is a published PGK1 inhibitor and has previously been shown to induce an accumulation of the reactive glycolytic metabolite methylglyoxal (MGO) in THP1 cells and other cell types^24,28^. Because CBR-470-2 itself does not contain a covalent reactive group, we hypothesized that CBR-470-2 likely acts by inhibiting its characterized cellular target PGK1. Knockdown of PGK1 via siRNA also recapitulated the NF-κB inhibitory effect of CBR-470-2 treatment, showing decreased induction of NF-κB target genes *IL1B* and *TNF* by qPCR (**Fig. 2D-F**). Collectively, these data suggested that CBR-470-2 inhibits NF-κB activation through a PGK1-dependent mechanism that might involve the accumulation of the reactive metabolite MGO.

**Figure 2.**
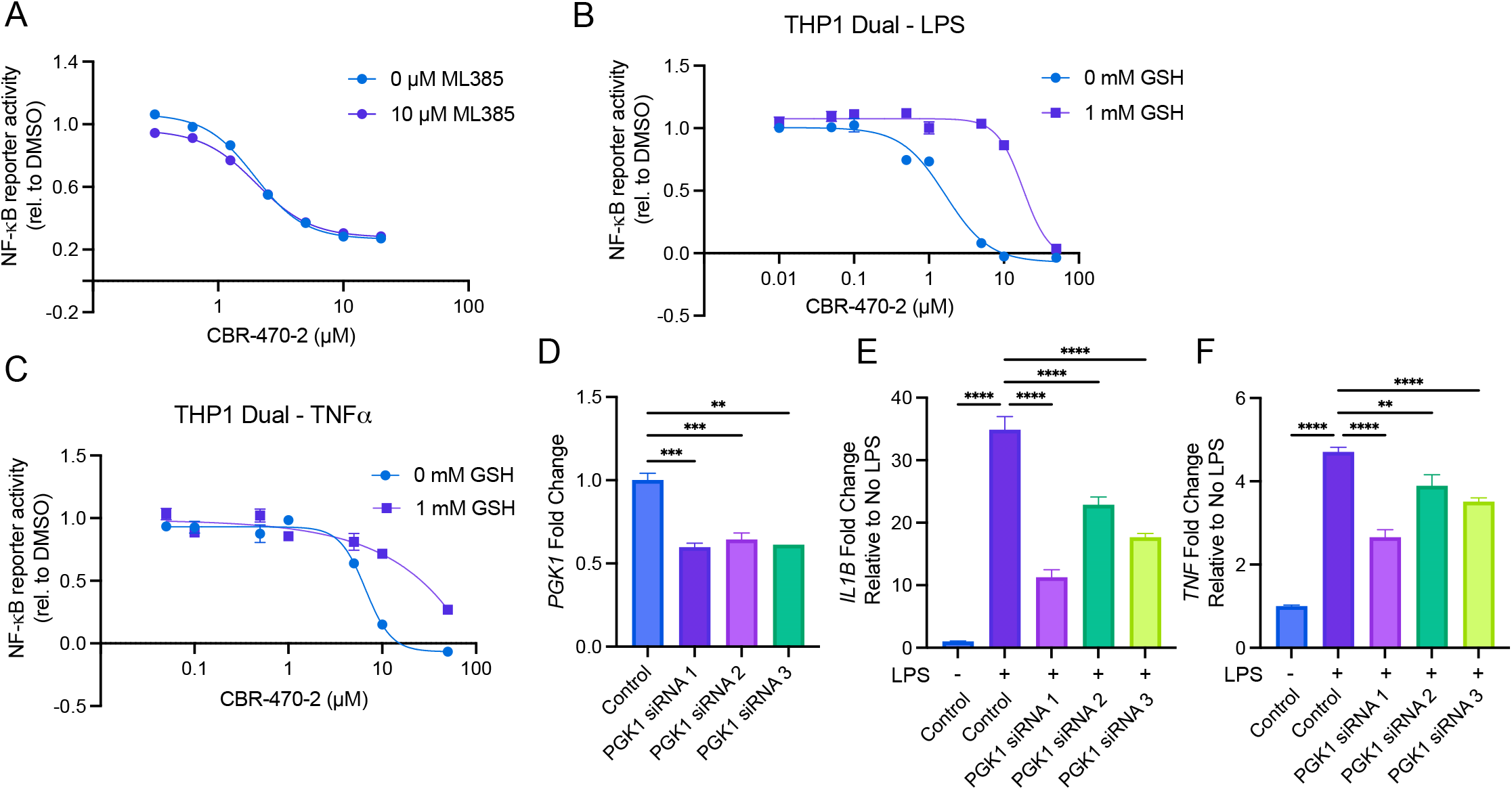
CBR-470-2 inhibits NF-κB through a mechanism involving PGK1 inhibition. (A) Relative NF-κB transcriptional activity as measured by SEAP reporter in THP-Dual cells cotreated with CBR-470-2 and 10 µM ML385 and then stimulated with LPS overnight. Error bars show SEM for n = 3 replicates. (B, C) Relative NF-κB transcriptional activity as measured by SEAP reporter in THP-Dual cells pretreated with 0 or 1 mM GSH for 30 min followed by CBR-470-2 for 6 h and then stimulated with LPS (B) or TNF-α (C) overnight. Error bars show SEM for n = 5 replicates. (D) Relative transcript level of *PGK1* 48 hr after knockdown with siRNA targeting *PGK1* in THP1 cells. Error bars show SEM for n = 3 replicates. Relative transcript level of *IL1B* (E) or *TNF* (F), from THP1 cells 48 hr following transfection with siRNAs targeting *PGK1* and then stimulated with LPS overnight, normalized to relative transcript level of respective untreated siRNA. Error bars show SEM for n = 3 replicates. **P<0.01, ***P<0.001, \*\*\*\**P*<0.0001 for ordinary one-way analysis of variance (ANOVA).

### Exogenous Methylglyoxal treatment recapitulates inhibition of NF-κB

We next sought to determine if increased levels of MGO were responsible for the NF-κB inhibitory effect of CBR-470-2. We found that exogenous treatment of MGO (1 mM, a concentration shown to mimic that of levels induced by CBR-470-2 in THP-1 cells^28^) could also inhibit activation of the NF-κB reporter by LPS- and TNF-α-induced stimulation in a dose dependent manner in both THP1 Dual cells and A549 Dual cells (**Fig. 3A-D**). Administration of MGO prior to TNF-α stimulation also resulted in the dose dependent reduction in nuclear localization of RelA, like CBR-470-2 treatment (**Fig. 3E**). Because MGO also directly modifies KEAP1 to activate NRF2 driven transcription, we additionally confirmed that inhibition of NRF2 by treatment with ML385 did not impact the inhibition of NF-κB observed by MGO in THP1 Dual cells stimulated with LPS (**Fig. 3F**). These data confirmed that exogenous methylglyoxal is capable of inhibiting NF-κB, supporting a model whereby MGO produced from PGK1 inhibition by CBR-470-2 is responsible for the compound’s inhibitory effect on NF-κB.

**Figure 3.**
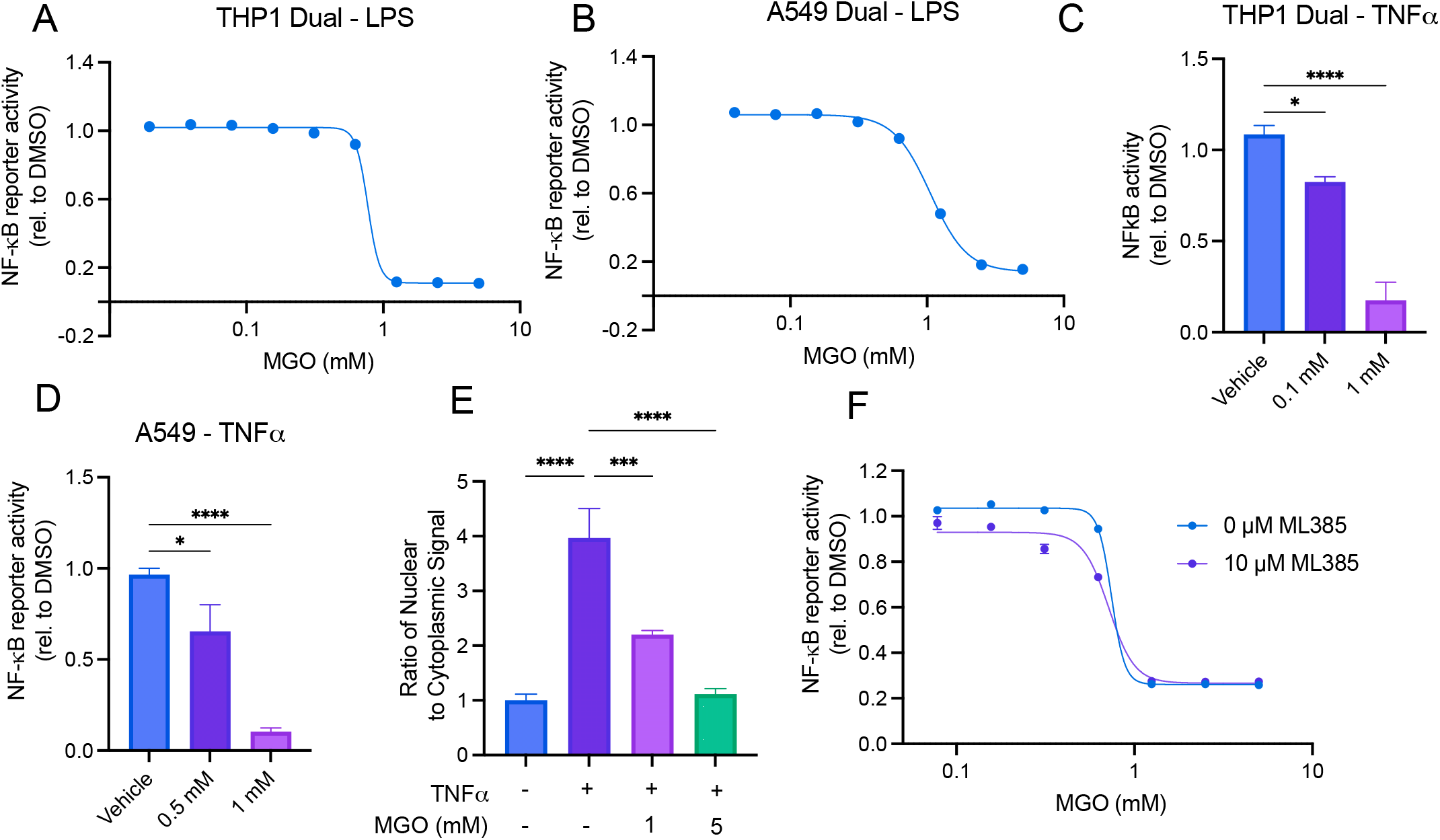
Exogenous methylglyoxal inhibits NF-κB. Relative NF-κB transcriptional activity as measured by SEAP reporter in THP Dual cells (A) or A549 Dual cells (B) pretreated with MGO for 6 h and then stimulated with LPS overnight. Error bars show SEM for n = 3 replicates. Relative NF-κB transcriptional activity as measured by SEAP reporter in THP Dual cells (C) or A549 Dual cells (D) pretreated with MGO for 6 h and then stimulated with TNF-α overnight. Error bars show SEM for n = 5 replicates. (E) Ratio of nuclear to cytosolic localization of RelA in A549 Dual cells pretreated with CBR-470-2 for 6 h and stimulated with TNF-α for 1 h. Error bars show SEM for n = 3 replicates. (A) Relative NF-κB transcriptional activity as measured by SEAP reporter in THP-Dual cells treated with the indicated concentrations of MGO in the presence or absence of ML385 (10 µM) and then stimulated with LPS overnight. Error bars show SEM for n = 3 replicates.

### CBR-470-2 Inhibits NF-κB through blocking phosphorylation of IKK proteins

We next sought to determine which step of the NF-κB signaling cascade is inhibited by increased levels of MGO. Previously, we demonstrated that MGO inhibits both KEAP1 and NLRP3 directly through the formation of non-enzymatically derived homo-dimeric protein crosslinks between cysteine and lysine, termed MICA modifications, which can be visualized as higher molecular weight species via target specific Western blotting. Accordingly, we evaluated if higher molecular weight species might form in response to either CBR-470-2 treatment or treatment with MGO. We observed that crosslinking was observed for all the major proteins in the NF-κB signaling cascade in response to both treatment conditions (**Fig. 4A-D**), as we observed higher molecular weight species of both the IκB kinases (IKKs), as well as the transcriptional components p50 and p65. To identify which of these crosslinking events, if any, might inhibit pathway signal relay, we monitored phosphorylation events in the pathway and degradation of IκBα via Western blotting. When cells were pre-treated with CBR-470-2 and then stimulated with TNF-a, we observed a marked decrease in the phosphorylation of both IKK kinases, as well as to p65 (**Figure 4E**). Relatedly, there was also a notable stabilization of IκBα levels at early timepoints, unlike the DMSO treated control, suggesting that signaling is not being relayed to this step of pathway activation. Although it appears that multiple steps in this signaling cascade are sensitive to modification by MGO, given that IKK phosphorylation is the most upstream step in this signaling cascade, we conclude that crosslinking of the IκB kinases (IKKs) is most likely is responsible for loss of signaling relay induced by CBR-470-2 treatment.

**Figure 4.**
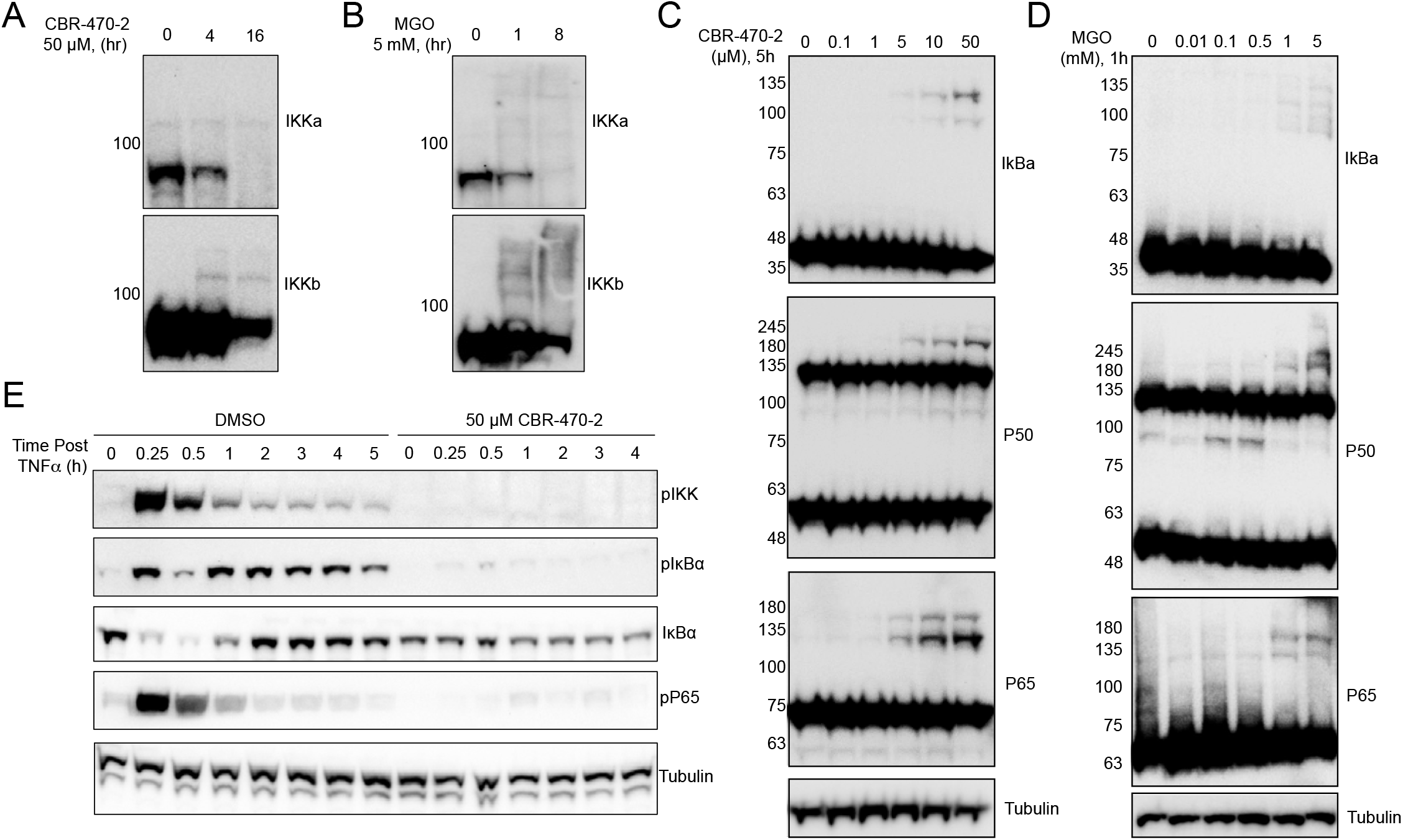
Accumulation of methylglyoxal promotes the crosslinking and inactivation of the NF-κB signaling complex. (A, B) Western blotting for IKKa and IKKb from THP-1 cells in response to CBR-470-2 (50 µM, A) or MGO (5 mM, B) at the indicated timepoints. (C, D) Western blotting for IkBa, P550, and p65 from THP-1 cells treated with the indicated concentrations of CBR-470-2 for 5 hours (C) or MGO for 1 hour (D). (E) Western blotting for phosphorylated levels of IKK, IκBa, or P65 after a 6-hour pre-treatment with 50 µM CBR-470-2 and then collection at the indicated timepoints after TNFa stimulation.

## Conclusion

Here, we have shown that attenuating the activity of the glycolytic enzyme PGK1, either pharmacologically with CBR-470-2 or by genetic depletion using siRNAs, results in the inhibition of the canonical NF-κB signaling pathway. As such, CBR-470-2 and its multiple reported analogs will likely serve as useful pharmacological tools for interrogating NF-κB signaling in cell-based systems. Future efforts to optimize the potency and to increase oral plasma exposure of this series will enable translation investigation into how PGK1 might be targeted therapeutically. Here, inhibition of NF-κB was found to arise from increased levels of MGO, a reactive metabolite that we have previously shown to inactivate both KEAP1 and NLRP3 through homo-dimeric protein crosslinks consisting of the non-enzymatically derived posttranslational modification MICA. Notably, we found that instead of promoting the crosslinking of one signaling protein, as was the case in these previous reports, MGO crosslinked all core signaling proteins evaluated, including p50, p65, IKKα, IKKβ, and IKBα, a result consistent with the previous observation that multiple NF-κB proteins possess reactive sensor cysteines that can be targeted by covalently acting electrophilic compounds. Given the preponderance of crosslinking events observed here, we suspect that likely many cysteine and arginine residues are modified by MGO, potentially highlighting a conserved capacity for sensing endogenous electrophilic species of this kind, as we have observed for other cellular electrophile sensors KEAP1 and NLRP3.

## Acknowledgments

This work was supported by the NIH (GM146865 to MJB; DK107604 and AG046495 to RLW). We thank Calibr at Scripps Research for providing CBR-470-2 and Luke Lairson for assistance with the high content imaging.

## Author Contributions

C.S., R.L.W., and M.J.B. designed research; C.S. and W.C. performed research; C.S. and W.C. analyzed data; and C.S., R.L.W., and M.J.B. wrote the paper.

## Methods

### Cell lines

WT THP1 cells, THP1-Dual cells, and A549-Dual cells were obtained from Invivogen and maintained according to the protocols of the manufacturer. HEK293T cells were obtained from ATCC and maintained in DMEM containing 10% fetal bovine serum (Gibco) and 1% penicillin-streptomycin (Gibco). All cells were maintained at 37°C in 5% CO2 in a humidified incubator.

### NF-κB Reporter Assay

THP1-Dual and A549 Dual cells were plated at a density of 20,000 cells per well in 30 µL of growth media in black clear bottom 384-well plates (Greiner). For assays with GSH, GSH was included in the initial plating media and incubated for 15 min prior to continuing protocol. MGO or CBR-470-2 was added in 10 µL of growth media to indicated final concentrations and incubated for 6 h. For assays with ML385 (10 µM), the compound was added concurrently to MGO or CBR-470-2. Following compound treatment, LPS (final concentration of 1 µg/mL) or TNF-α (final concentration 0.1 ng/mL) was added in 10 µL of media and incubated overnight. The next morning, 30 µL of Quanti-Blue (Invivogen) was added to each well and incubated at 37 °C until a visible absorbance signal was obtained at 655 nm.

### RT-qPCR

Cells were treated for 6 h with MGO or CBR-470-2 before treatment with LPS (1 µg/mL) or TNF-α (1 ng/µL) overnight. For assays involving GSH treatment, GSH was supplemented to the initial plating media and incubated 15 min prior to continuing the protocol. Cells were pelleted and lysed following the RNeasy Mini Kit manufacturer’s protocol to isolate total RNA. Using the High-Capacity cDNA Reverse Transcription Kit, 400 µg of RNA was converted to cDNA. Each reaction was then then diluted 1:4 with DNase/ RNAse free water. qPCR reaction mixtures were prepared with Power SYBR Green PCR Master Mix and gene specific primers as noted below. RT-qPCR reactions were performed using the QuantStudio™ 7 Flex with an initial melting period of 95 °C for 10 min and then 40 cycles of 15 s at 95 °C, 1 min at 60 °C.

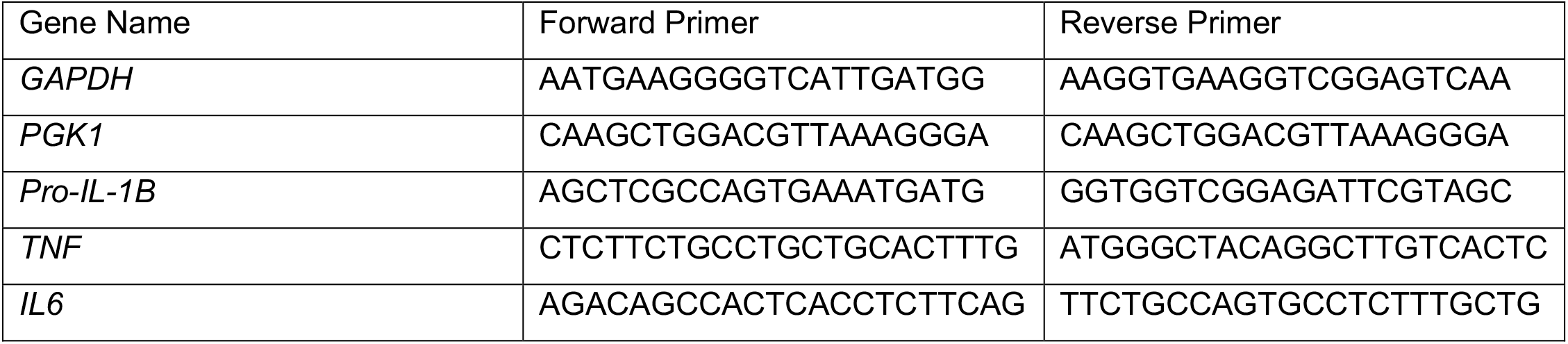

### siRNA

PGK1 knockdown using siRNA was performed using Lipofectamine™ RNAiMAX Transfection Reagent following the manufacturer’s protocol with 9 µL of RNAiMAX reagent and 3 µL of 10 µM siRNA in 300 µL of Optimem per 2 million cells in a 6 well tissue culture dish. 2 days after transfection, cells from the siRNA experiments were treated with LPS (1 µg/mL) overnight and then harvested for gene knockdown confirmation and qPCR of NF-κB target transcripts.

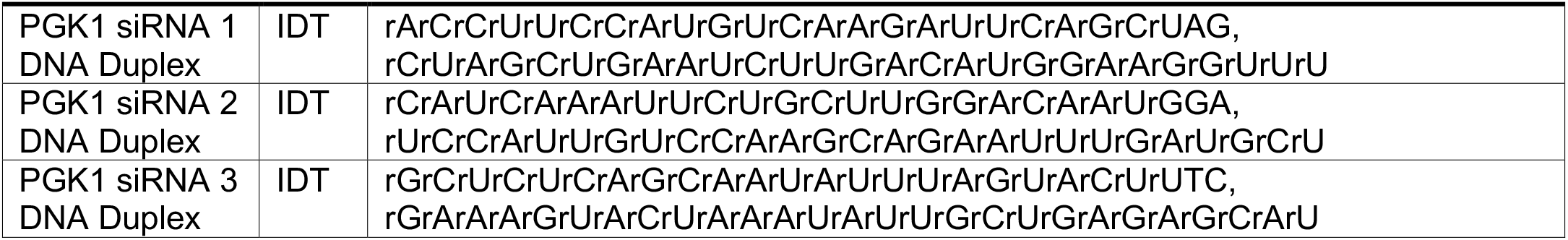

### Immunofluorescence

A549 Dual Reporter cells were plated in poly-D-lysine coated 96 or 12 well plates at a density of 30,000 or 300,000 cells per well respectively and allowed to adhere overnight. The next day, cells were treated with CBR-470-2 or MGO for 6 h and then NF-κB stimulated by treatment with TNF-α (1 ng/µL) for 1 h. Cells were fixed with 4% paraformaldehyde for 15 min at room temp and then washed four times with PBS. Cells were incubated with anti-NFkappaB p65 (1;200, sc-8008, Santa Cruz Biotechnology) and anti-mouse Alexa Fluor 488 conjugated antibodies (1:200, Thermo Fisher) overnight at 4°C. The following day, cells were washed three times with PBS and incubated for 30 min with 5 µg/mL Hoechst before imaging. Cells in 96 well format were imaged on the Cellinsight CX5 HCS Platform and the nuclear to cytoplasmic NF-κB P65 ratio was quantified using the intensity function of the Cellinsight High Content Analysis Platform.

### Immunoblotting and Immunoprecipitation

WT THP1 cells were treated with CBR-470-2 or MGO for the indicated time periods with or without stimulation by TNF-α. Cells were pelleted via centrifugation and lysed in RIPA buffer with 100X phosphatase/protease inhibitor (Cell Signaling, #5872) via probe sonication. Insoluble material was separated via centrifugation and protein concentration in soluble lysate was quantified on a nanodrop instrument. Protein concentrations were normalized and 4X SDS loading dye containing 10% beta mercaptoethanol was added to the samples. Approximately 50 µg of protein per sample was loaded per well for separation via SDS-PAGE (SDS– polyacrylamide gel electrophoresis) in 12, 15, or 17 well gels (4-20% Bis-tris BOLT gels, Invitrogen) with 1X MOPS Buffer (Invitrogen) at 200 V for 40 min. Protein was transferred to PVDF (polyvinylidene difluoride) membranes using semi-dry transfer at 20 V for 1 h, and then blocked in 5% blotting grade milk in TBST (20 mM Tris, 137 mM NaCl, 0.1% Tween 20) for 1 hr. Each membrane was then incubated overnight in respective primary antibody in 5% BSA or 5% blotting grade milk at 4°C. The following day, blots were washed six times in TBST over 30 min, incubated with species specific secondary antibody (LI-COR, 1:5000), (HRP, 1:3333) in 5% milk for 1 h, and then washed 10X in TBST over 1 h. HRP blots were incubated in HRP substrate (West Dura) and then imaged on a ChemiDoc Imaging system.

| Protein | Supplier | Catalog Number | Host Species | Dilution |
| --- | --- | --- | --- | --- |
| Tubulin | Sigma | T6557 | Mouse | 1:2000 (BSA) |
| IKKα (3G12) | Cell Signaling | 11930 | Mouse | 1:1000 (BSA) |
| IKKβ (D30C6) | Cell Signaling | 8943 | Rabbit | 1:1000 (BSA) |
| Phospho-IKKα/$\beta$ (Ser176/180) (16A6) | Cell Signaling | 2697 | Rabbit | 1:1000 (BSA) |
| Phospho-NF-κB p65 (Ser536) (93H1) | Cell Signaling | 3033 | Rabbit | 1:1000 (BSA) |
| IκBα (L35A5) | Cell Signaling | 4814 | Mouse | 1:1000 (BSA) |
| Phospho-IκBα (Ser32) (14D4) | Cell Signaling | 2859 | Rabbit | 1:1000 (BSA) |
| NF-κB p65 (D14E12)<br>XP® | Cell Signaling | 8242 | Rabbit | 1:1000 (BSA) |
| IKKγ | Cell Signaling | 2685 | Rabbit | 1:1000 (BSA) |
| FLAG | Sigma | F1804 | Mouse | 1:1000 (Milk) |
| IL-1β | GeneTex | GTX74034 | Rabbit | 1:1000 (BSA) |

---

**Supplementary Figure 1.**
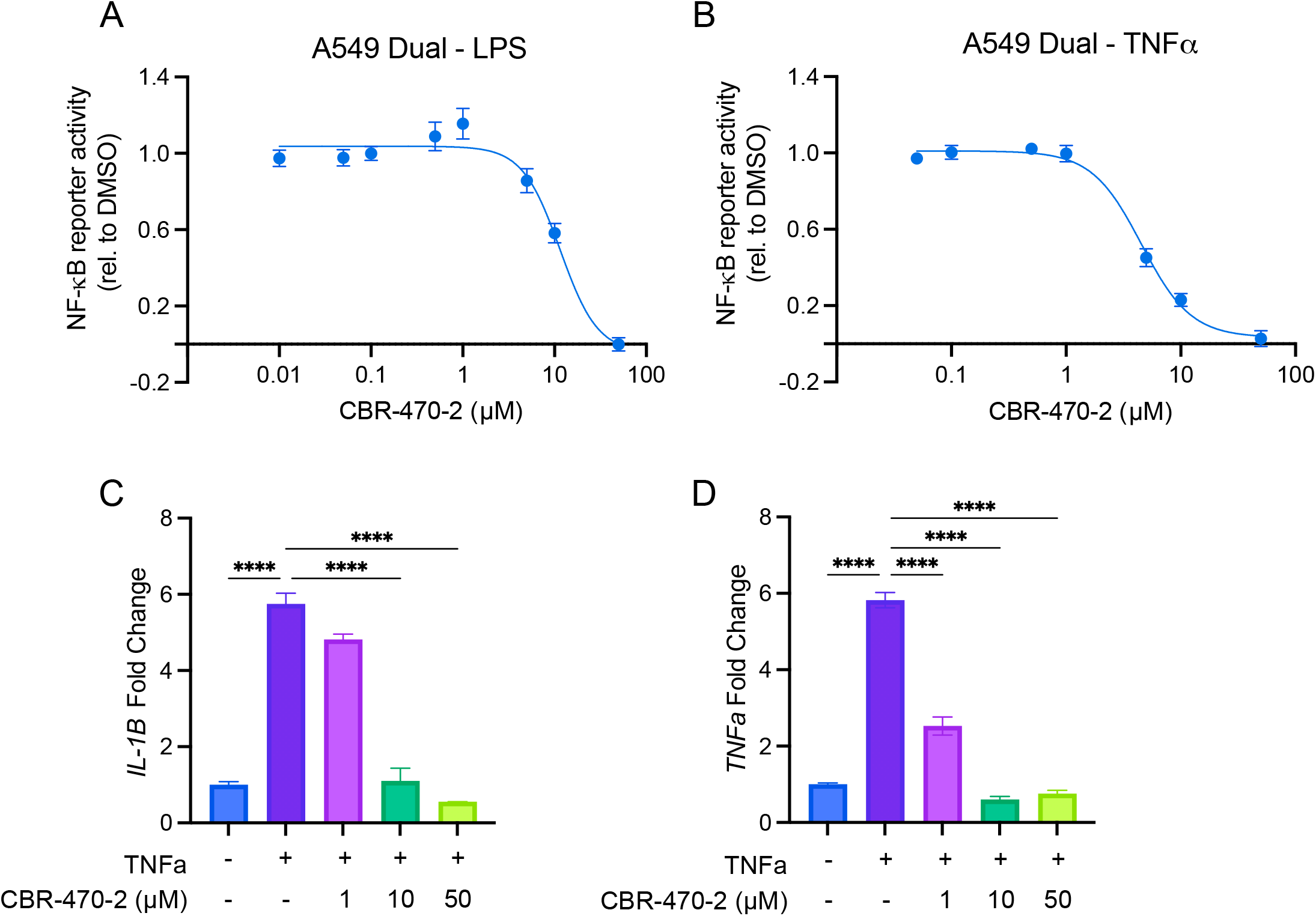
CBR-470-2 inhibits NF-κB in multiple cell types and multiple NF-κB stimuli. (A,B) Relative induction of NF-κB as measured by SEAP reporter in A549-Dual cells pretreated with CBR-470-2 for 6 h and then stimulated with LPS (A) or TNF-α (B) overnight. Error bars show SEM for n = 5 replicates (A) or n = 4 replicates (B). (C, D) Relative transcript level of *IL1B* (C) or *TNF* (D) from THP1 cells pretreated with CBR-470-2 for 6 h and then stimulated with TNF-α overnight. Error bars show SEM for n = 3 replicates. \*\*\*\**P*<0.0001 for ordinary one-way analysis of variance (ANOVA) with Tukey correction for multiple comparisons between conditions.

**Supplementary Figure 2.**
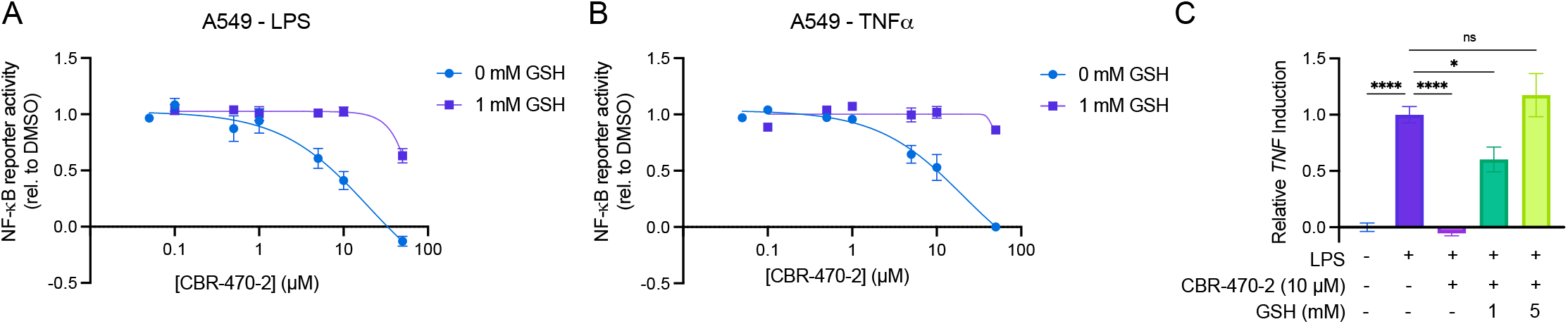
CBR-470-2 inhibitory activity is quenched by exogenous GSH. (A, B) Relative induction of NF-κB as measured by SEAP reporter in A549-Dual cells pretreated with 0 or 1 mM GSH for 30 min followed by CBR-470-2 for 6 h and then stimulated with LPS (A) or TNF-α (B) overnight. Error bars show SEM for n = 4 replicates. (C) Relative transcript level of *TNF* from THP1 cells pretreated 0, 1, or 5 mM GSH for 30 min followed by 10 µM CBR-470-2 for 6 h and then stimulated with LPS overnight. Error bars show SEM for n = 3 replicates. *P<0.05, \*\*\*\**P*<0.0001 for ordinary one-way analysis of variance (ANOVA) with Dunnett correction for multiple comparisons between conditions.

